# Ice thickness control and measurement in the VitroJet for time-efficient single particle structure determination

**DOI:** 10.1101/2023.10.09.561488

**Authors:** Rene J.M. Henderikx, Maaike J.G. Schotman, Saba Shahzad, Simon A. Fromm, Daniel Mann, Julian Hennies, Thomas V. Heidler, Dariush Ashtiani, Wim J.H. Hagen, Roger J.M. Jeurissen, Simone Mattei, Peter J. Peters, Carsten Sachse, Bart W.A.M.M. Beulen

**Author notes:** These authors contributed equally (Rene J.M. Henderikx, Bart W.A.M.M. Beulen).

## Abstract

Embedding biomolecules in vitreous ice of optimal thickness is critical for structure determination by cryo-electron microscopy. Ice thickness assessment and selection of suitable holes for data collection are currently part of time-consuming preparatory routines performed on expensive electron microscopes. To address this challenge, a routine has been developed to measure ice thickness during sample preparation using an optical camera integrated in the VitroJet. This method allows to estimate the ice thickness with an error below ± 20 nm for ice layers in the range of 0 – 70 nm. Additionally, we implemented pin printing to reproduce and control sample deposition in the VitroJet. The median ice thickness can be reproduced with a standard deviation below ± 11 nm for thicknesses up to 75 nm. Therefore, the ice thickness of buffer-suspended holes on an EM grid can be tuned and measured within the working range relevant for single particle cryo-EM. Single particle structures of apoferritin were determined at two distinct thicknesses of 30 nm and 70 nm. These reconstructions demonstrate the importance of ice thickness for time-efficient cryo-EM structure determination.

**Highlights:**

- Methods in the VitroJet allow for on-the-fly ice thickness tuning and measurement
- The optical camera can estimate ice thickness ranging from 0 – 70 ± 20 nm
- Pin printing enables to reproduce and control median ice thickness up to 75 ± 11 nm
- Structures of apoferritin require 3.7 x fewer particles in 30 compared to 70 nm ice

## 1. Introduction

Over the past decade, cryogenic electron microscopy (cryo-EM) has experienced remarkable advancements in its ability to elucidate the structures of biological macromolecules ^1,2^. The ability to resolve large protein complexes as well as smaller membrane proteins makes it frequently the method of choice for structural biologists ^3^. However, to refine atomic models and accurately dock ligands for structure-based drug design, the reconstructed EM-maps require resolutions better than 3 Å^4,5^. In the current cryo-EM workflows, it is common to do multiple iterations of microscope screening to evaluate sample quality, which remain a time-consuming practice ^6^. Apart from the molecular particle properties, such as size, structural order and symmetry, ice thickness is a key determinant for the achievable resolution ^7,8^. Ideally, the ice thickness is such that only a monolayer of particles fit into the ice layer, ensuring full particle hydration and an unobstructed view of the particle in the micrograph. Therefore, the desired ice thickness is dependent on particle dimensions. When the deposited layer is too thin, the particles are not sufficiently hydrated and often pushed out of the thin regions, while thicker layers add additional noise from the vitreous ice and allow for undesired particle overlap ^9^. When collecting data on the protein aldolase with an ideal ice thicknesses of 10 – 20 nm, less than 400 movies were shown to be sufficient to obtain a resolution of 2.8 Å^10^. With the current state of the art cryo-electron microscopes, a data collection of 400 movies only requires around one hour of beam time ^11^. Despite the key role of ice thickness for structure determination, traditional grid preparation methods lack feedback and control over layer thickness.

When using conventional grid preparation, the grids quality can only be assessed once they are loaded and imaged inside the electron microscope. To decrease the time required for the identification of optimal preparation conditions, several studies focused on ice thickness determination by cryo-EM imaging. The principle of this method relies on comparing the intensity of foil holes in presence and absence of an energy filter using the mean free path of the inelastically scattered electrons ^12,13^. The estimation of ice thickness was demonstrated for different microscope setups and grid types ^14–16^. Although these methods help in identifying the optimal grid preparation conditions and targeting holes with suitable ice thickness, the procedure requires additional beam and operator time that is commonly limited. A few methods that do not involve electron microscopy were put forward to estimate layer thickness based on interference of visible light, either in reflection or transmission. Most methods measured optical intensity as a proxy for ice thickness where no mathematical correlation was presented ^17–19^. Nonetheless, two studies showed quantification of ice thickness based on the use of neural networks ^20,21^. To increase throughput and cost-efficiency, determination of layer thickness during grid preparation would be of great benefit ^22^.

Classical grid preparation methods for sample deposition rely on pipetting a droplet of a few microliters onto a grid and subsequently wicking away the excess liquid through blotting paper, leaving a thin enough layer for the penetration of the electrons. As this method does not provide accurate control over the liquid removal, variations in thickness from one grid to another and within one grid commonly occur. Consequently, multiple grids need to be screened before finding ice thickness optimal for data acquisition ^23^. Alternatives to blotting as a means for sample preparation have been proposed, which can be classified into droplet-spraying and scribing methods. In the droplet-spraying methods, one generates droplets that are sprayed onto the perforated foil of the grid ^24–26^. Afterwards, the droplets spread, and in some cases excess sample is wicked away by, for example, nanowires ^27^. For scribing methods, a tip moves over the grid and leaves a thin layer of liquid behind ^19,28^. The deposition of an optimum sample thickness is experimentally dependent on sample fluid and grid properties ^29,30^. Increasing reproducibility and enhancing control over ice thickness would yield a significant improvement in the workflow ^31^.

To obtain the desired layer thickness in grid preparation, we present a routine that estimates the layer thickness in individual holes of the perforated foil from optical images. A camera with on-axis illumination captures images of the grid before and after sample deposition prior to vitrification. The intensity inside the holes is used to determine the expected ice thickness, which is in agreement with an optical model based on a thin layer interferometry. Additionally, we demonstrate how ice thickness can be reproduced and controlled using pin-printing as a technique for sample deposition. In this work, the parameters of writing velocity and the standoff distance are shown to experimentally correlate with the ice thickness, which are supported by fluid dynamics scaling. Moreover, we demonstrate that targeted data collection based on ice thickness is beneficial for structure determination of apoferritin. As a result, sample deposition by pin printing and the obtained optical thickness measurement can minimize the necessary cryo-EM beam time to achieve optimal sample thickness for protein structure determination.

## 2. Results

### 2.1. Experimental setup

#### 2.1.1. Grid preparation

To tackle the challenges associated with optimizing ice thickness in the cryo-EM workflow, we set out to characterize the capabilities of the VitroJet machine during grid preparation ^32^. Sample deposition occurs in a temperature- and humidity-controlled climate chamber, and an integrated camera records the process (Fig. 1A, details in Materials and Methods). The light source emits white light, passing through an absorptive and long-pass filter to make the beam uniform and match the transmission range of the objective lens. Using two mirrors and an objective lens, the light is directed to the autogrid where it illuminates the area of interest. The reflected light is captured by the objective, and through another mirror imaged by the camera. For sample deposition itself, a sample of interest is loaded in an automated pipette and inserted in the climate chamber. A solid pin dips into the pipette tip to pick up the sample and to write it onto the autogrid by pin-printing ^28^. Both pin and the autogrid are temperature controlled with respect to dewpoint temperature in the climate chamber to minimize the evaporation of the thin layer of the sample.

**Figure 1:**
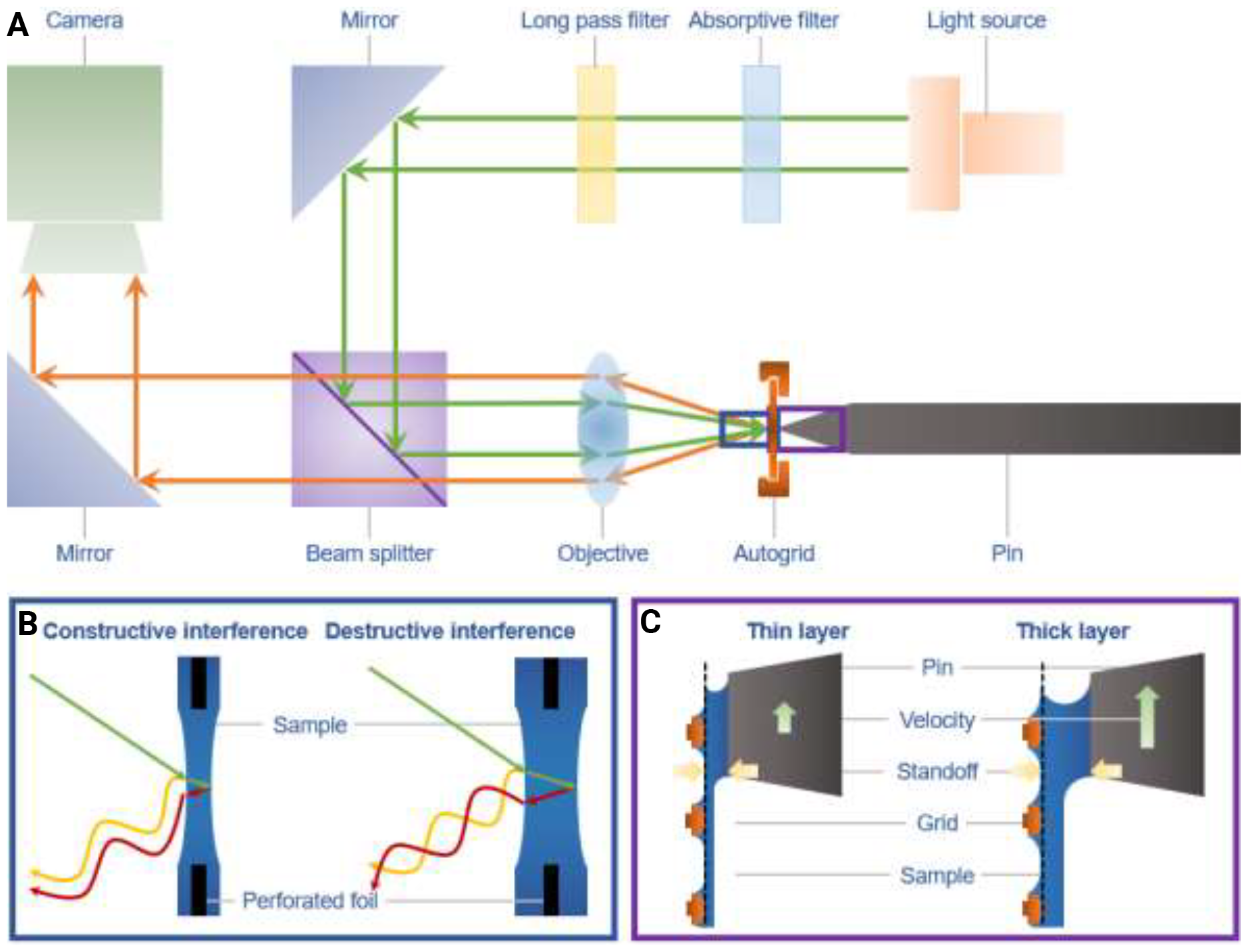
Schematic overview of the deposition and imaging setup integrated in the VitroJet. (A) White light is generated and passes through filters, a mirror, beam splitter and an objective lens. The light is reflected at the sample and autogrid, passes through the objective again, the beam splitter and another mirror, and finally arriving at the camera. A sample is deposited onto the autogrid using a solid pin that moves laterally with respect to the grid. (B) Zoomed in sections for the thin and thick sample layers indicating how fractions of the light reflect on the first and second air-water interface before vitrification. Depending on the layer thickness, the reflected light can yield constructive or destructive interference. (C) Magnified view of the sample deposition by pin printing showing how sample thickness depends on the writing velocity and standoff distance. Decreasing writing velocity or standoff distance results in a thinner layer remaining on the grid.

The camera records images with a field of view of 1.0 x 0.8 mm, where the intensity is dependent on the sample thickness (Fig. 1B). Illuminated light is transmitted or reflected at both the front and back air-water interface of the sample. This results in a phase shift between the reflected light wave from the different interfaces, leading to a change in total intensity. As the layer thickness increases, the reflected intensity will increase until a quarter wavelength of the light is reached, at which constructive interference occurs. Above this thickness, the intensity decreases again until destructive interference occurs, and the Newton interference rings form.

During pin printing, the sample bridges the gap between the pin and the grid. Since both are hydrophilic, the sample tends to remain in contact with the pin and follow its movement. When increasing the velocity, the sample has more difficulty to keep up with the pin motion and therefore, a thicker layer remains on the grid. Similarly, a larger standoff distance results in the deposition of a thicker sample layer (Fig. 1C). Sample deposition started in the center of the grid, and spiraled outwards to cover a circular area with a diameter of approximately 800 µm.

#### 2.1.2. Cryo-EM ice thickness determination

After loading the grids in the electron microscope, an atlas is recorded from the region of interest with vitreous ice layer (Fig. 2A). The deposited sample is located in the center of the grid, significantly speeding up atlas acquisition since only 9 tiles instead of the regular 35 are recorded. The grid area outside the deposited area has a speckled appearance due to ice contamination on the perforated foil. The region of the deposited area appears uniformly grey, indicating a clean area with ice suitable for analysis.

**Figure 2:**
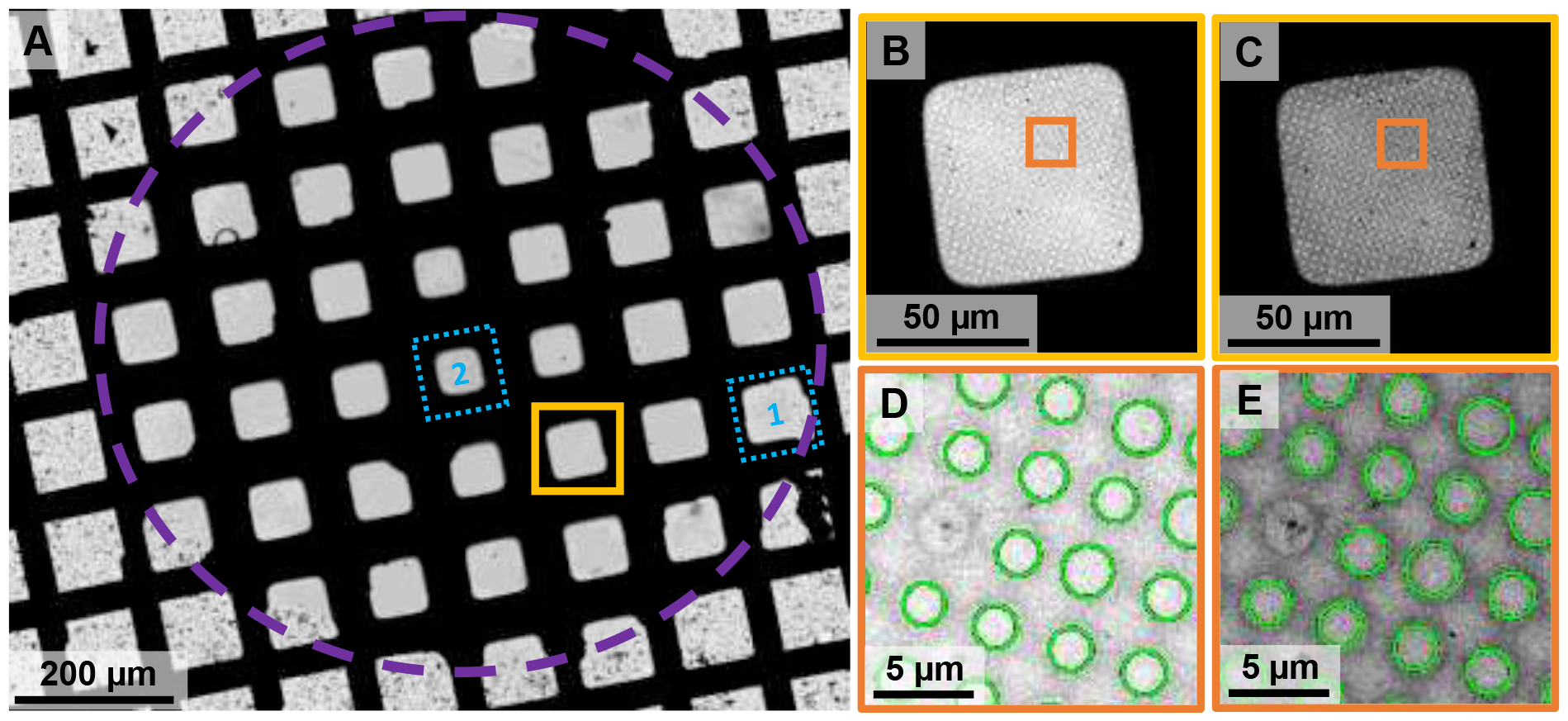
Cryo-EM images after vitrification (A) The low magnification atlas of the grid, showing the spiral deposition pattern in the center of the image, indicated with a dashed purple circle. Data collected for a 3D reconstruction was performed in two squares marked with a dashed blue line. Image of the square highlighted in yellow in the atlas (B) without energy filter and (C) with the energy filter present. Note the increased contrast in the square with the energy filter. (D and E) Enlarged view of the orange area in the square image, indicating the holes used for the analysis in green.

The ice thickness from cryo-EM images is estimated using the energy filter method. For each square, medium magnification images in the absence (Fig. 2B) and presence (Fig. 2C) of an energy filter are recorded at the same location. When the energy filter is inserted, electrons that are inelastically scattered in the sample are removed and do not contribute to the filtered image. The resulting difference in contrast with respect to the non-filtered image can be used to assess ice thickness.

For each square image in absence and presence of the energy filter, the region close to the grid bars of the square appears as a dark ring and contains thick ice. This is excluded from further analysis using intensity thresholding, and homogeneously grey area within the squares is used for processing. The holes in the perforated foil are detected, and subsequently the average intensity in the center of each hole is determined (Fig. 2DE).

The intensity in both the unfiltered (*I*_*tot*_) and the filtered (*I*_*zlp*_) image is related to the ice thickness (*t*), as the number of electrons that inelastically scatter in the sample depends on its thickness ^12^. Using the apparent mean free path of the electrons (*λ*_*inel*_) and the ratio of intensities, the ice thickness can be determined ^14^. To ensure that empty holes correspond to a thickness of zero nanometer, a correction factor (*C*) was introduced ^15^.

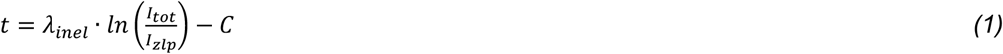

Both the mean free path and correction factor can be influenced by the used microscope settings. The mean free path is dependent on objective aperture size and energy filter slit width. The offset for the correction factor could originate from delocalization due to the applied defocus during imaging. In our experiments, two different microscopes were used to determine the ice thicknesses for correlation to optical intensity and pin print parameters. The correction factor is determined by shifting the peak of the empty holes in the histogram to zero resulting in respective correction factors of 9 nm and 10 nm. For the intensity correlation, a mean free path of 485 nm was used corresponding to a 200 kV microscope and medium magnification during imaging ^15^. For the pin-print correlation, the mean free path for the 300 kV microscope was calibrated to 352 nm using tomography (Supplementary Materials A.1). This resulted in a root mean square deviation of 6 nm for the energy filter method to determine the ice thickness.

### 2.2. Optical ice thickness measurement

#### 2.2.1. Intensity determination

We exemplify the same grid from the electron microscope and compare it with the optical images recorded during the deposition process. Images of the grid before and after deposition are obtained by the camera. The image before deposition shows the mesh of the grid and its perforated foil, including a slight variation in illumination across the field of view (Fig. 3A). Due to the reflection, the copper mesh appears bright in the image, while the perforated carbon foil looks darker. The image after deposition shows the grid just before vitrification, where the grey level in the wetted area changed in comparison with the dry grid (Fig. 3B). Furthermore, the sample accumulates at the hydrophilic grid bars due to wicking by capillary forces, leading to a very thick layer that appears as dark surroundings around each square. Conversely, the central regions of the squares appear homogeneously bright. The zoomed-in images show that the holes in the foil are clearly detectable (Fig. 3C), including intensity changes originating from the deposited sample (Fig. 3D). The difference in intensity depends on layer thickness due to thin film interferometry.

**Figure 3:**
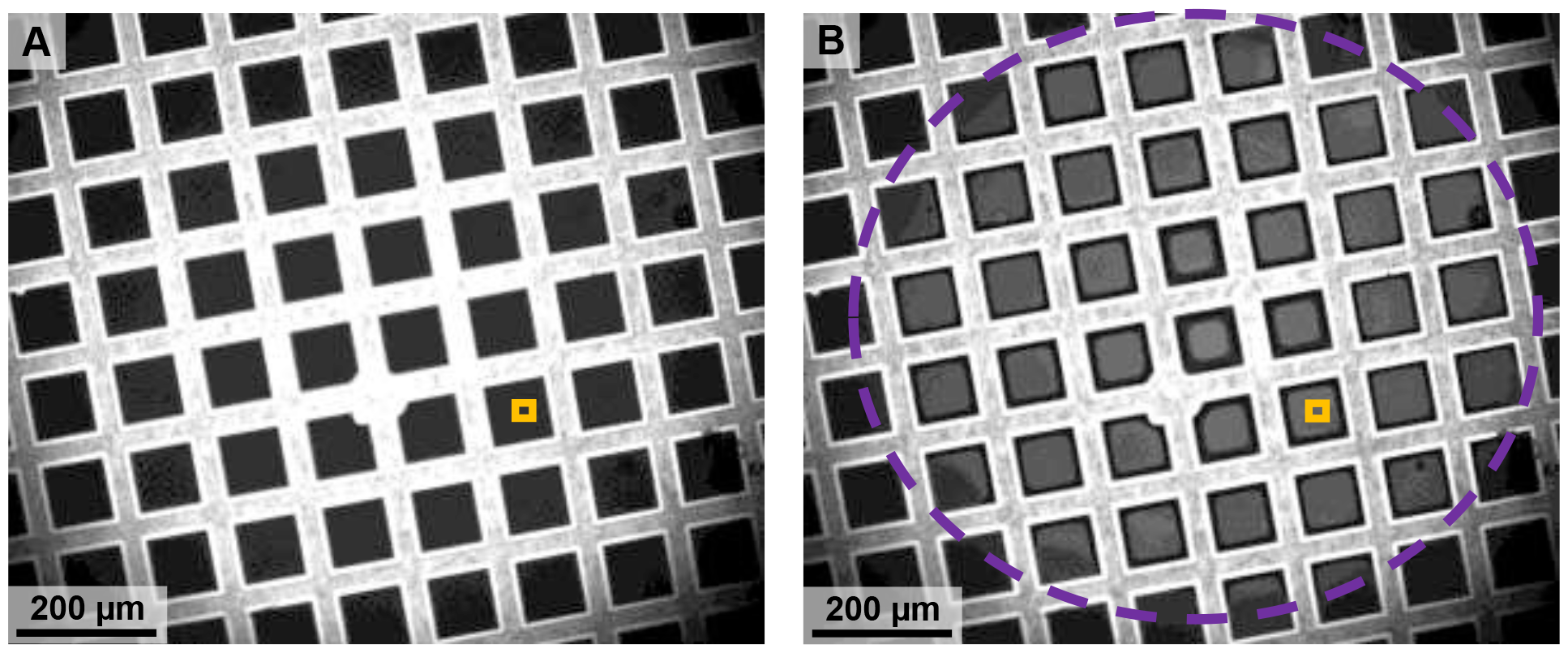
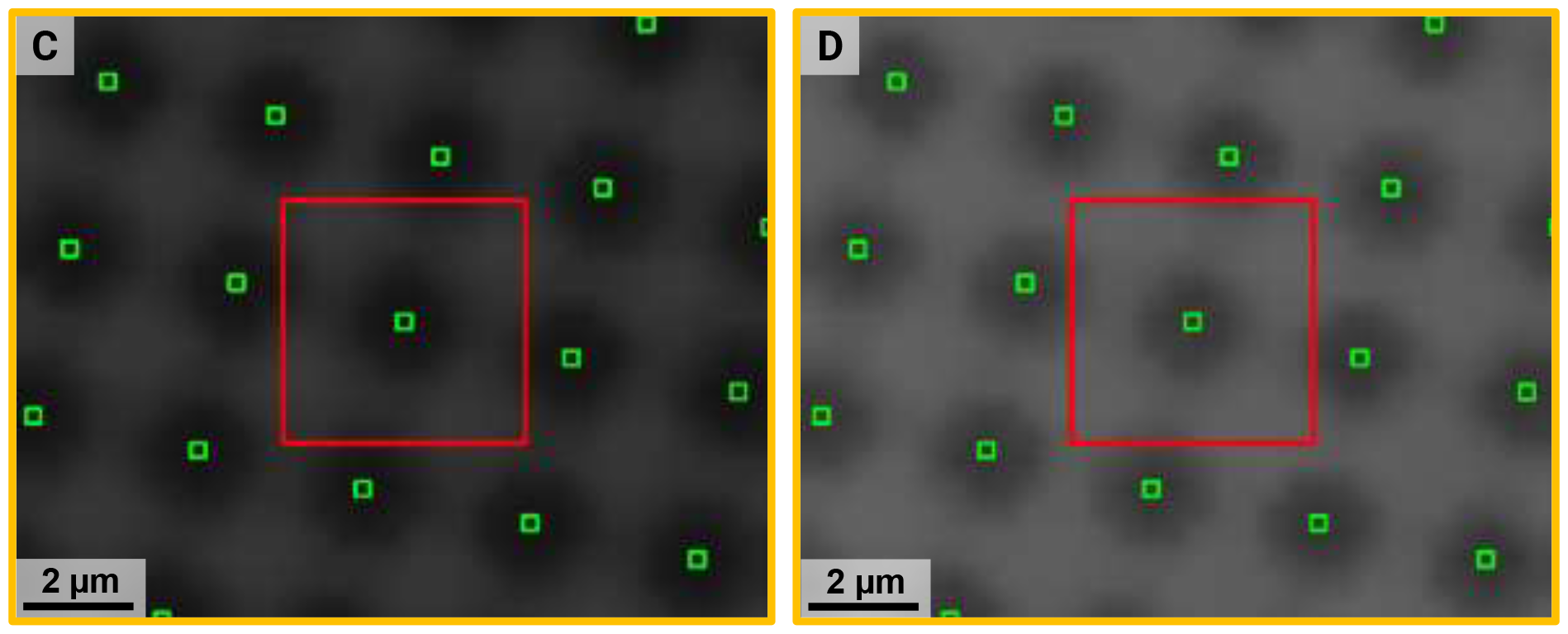
Images acquired by the VitroJet camera before and after sample deposition, prior to vitrification of the same grid shown in Figure 2. (A) Image showing the 200 mesh R2/1 grid including the copper mesh and the perforated carbon foil before deposition. (B) Image of the same grid after deposition, where the sample changes the reflected light intensity. The purple dashed circle highlights the area in which deposition by pin printing occurred. (C and D) Magnified image with bilinear interpolation of A and B, respectively, where the individual holes of the carbon foil are visible. The area used for intensity determination is marked in green, and the region to detect local surrounding carbon intensity in red.

For quantitative image analysis, the principal grid features can be detected by thresholding. First, grid squares are identified based on the brightness and known mesh dimensions. Second, for each square, the dark area close to the grid bars due to wicking are excluded from further analysis by intensity thresholding. In the remaining area within each square, holes are detected using the Hough transform ^33^. The circle detection results in an initial estimation of the location of the hole center. Due to the small hole size and corresponding limited number of pixels per hole, the center of the holes needs to be refined with sub-pixel accuracy. Therefore, the image is blurred with a kernel of 3 x 3 pixels, and the area around the initial hole location is enlarged ten times using bilinear interpolation. The hole center is determined by the centroid of the patch of pixels that lies within a range of 30% from the minimum intensity value in the enlarged window. The hole intensity is taken as the mean intensity of a window of 5 x 5 pixels around the center.

Although there are multiple pixels in each detected hole, an empty hole in the image still shows some intensity due to diffraction from the carbon foil and imperfections in the optical system. To compensate for this background signal, the difference between the intensity of each hole before 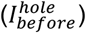 and after deposition 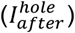 was used.

Additionally, the intensity of the light source across the grid is not completely uniform. To correct for this effect, the maximum intensity of the carbon surrounding each individual hole is locally determined 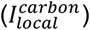. Subsequently, this value is normalized using the maximum carbon intensity in the total field of view 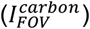. In summary, the background corrected intensity for each hole (*I*^*hole*^) is calculated by performing a compensation for the illumination and diffraction according to:

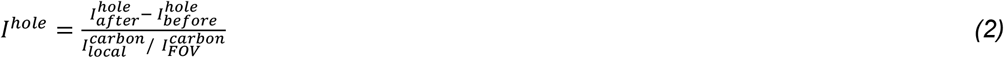

#### 2.2.2. Spatial correlation

To validate the spatial correlation between optical camera intensity and cryo-EM ice thickness, we inspected the respective grid maps. The background corrected intensity colormap is plotted over the optical image (Fig. 4A), and the ice thickness calculated by the energy filter method superimposed with the cryo-EM atlas (Fig. 4B).

**Figure 4:**
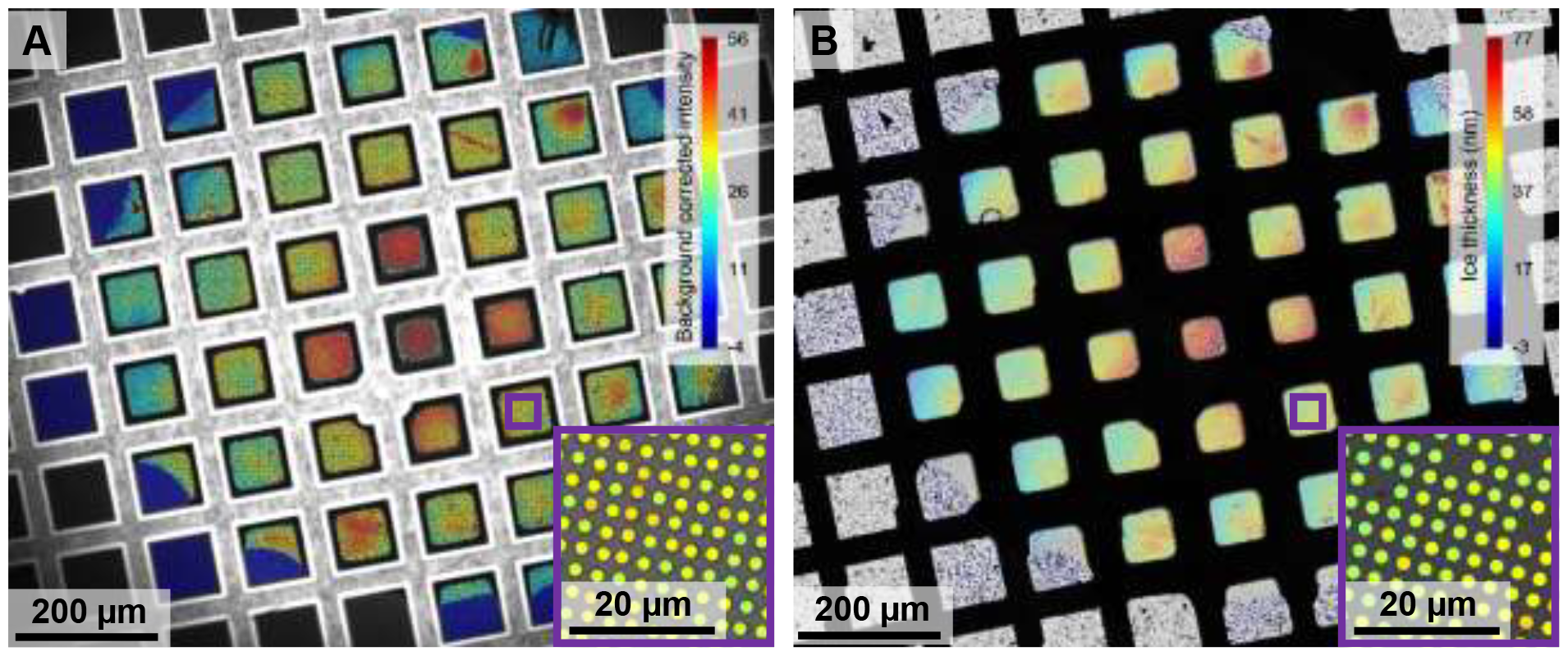
Color overlay of the background corrected intensity from the optical camera, and of the ice thickness measured in the electron microscope on the atlas. (A) Background corrected intensity overlaid over the optical image of the camera in a color plot. (B) Ice thickness with color plot superimposed on cryo-EM atlas obtained in the electron microscope. The matching colors reveal a qualitative correlation between background corrected intensity and ice thickness.

The holes with lower intensity and ice thickness are colored blue, while the holes of bright and thicker ice are colored red. This way the spatial distributions of the background corrected intensity and that of the ice thickness visually correlate very well. Dark blue colors outside the printing area show that no sample was deposited in these squares. In the deposited region of both color maps, a gradient can be discerned emanating from center of the grid to the outside rim. Hence, this observation reveals that optical images may be sufficient for judging optimal ice thickness replacing the need for energy filtered images.

#### 2.2.3. Intensity – thickness correlation

To evaluate the relationship between ice thickness and background corrected intensity from the optical camera quantitatively, a total of 18 grids are analyzed with the test sample apoferritin. To compare holes from the two imaging methods, images must be brought in register using a rigid transformation. After subsequent hole identification and assignment, a total number of 101,673 unique holes with both optical intensity and ice thickness became available and were plotted (Fig. 5A). Two reduced one-dimensional histograms of the data reveal two continuous distributions with most holes corresponding to thinner ice (20 - 50 nm), while there are fewer holes available at larger thickness (Fig. 5BC). An additional peak is found around zero, corresponding to empty holes.

**Figure 5:**
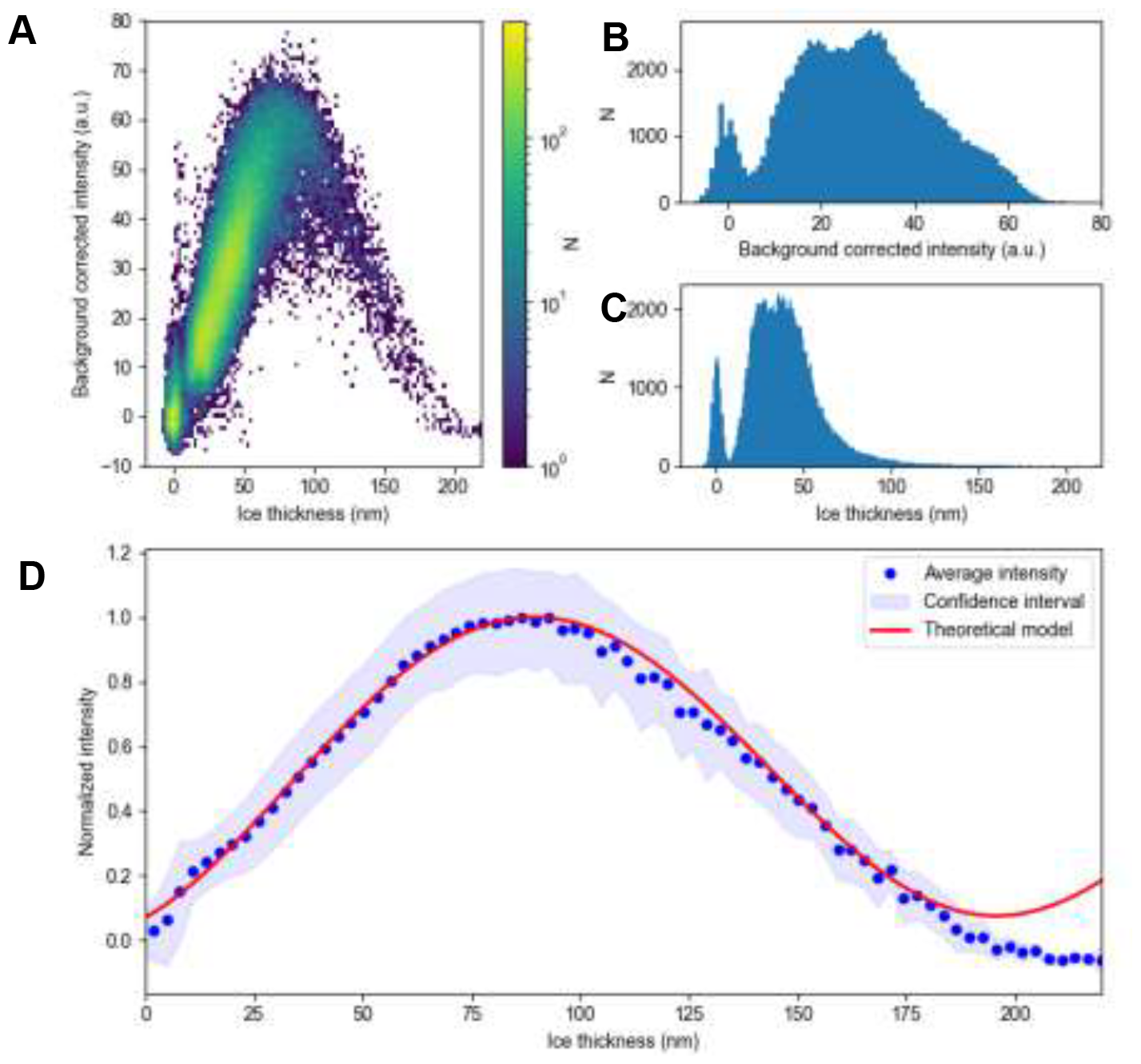
Relationship between optical intensity of VitroJet camera and ice thickness in the microscope. (A) Scatterplot of identified 101,673 holes with the optical intensity and corresponding ice thickness estimated by the energy filter method, where the number of occurrences for each pixel is indicated by color. Reduced one-dimensional histograms of (B) background corrected intensity and (C) ice thickness. (D) Average background corrected intensity after normalization by the maximum intensity in blue, in combination with the standard deviation presented as confidence intervals. Fit of the optical model in red, demonstrating that the expected reflectance based on thin film interferometry matches the experimental data in the 0 – 180 nm interval.

Afterwards, the data pairs are divided into bins of 3 nanometer based on thickness. For each bin, the average and standard deviation of the background corrected intensity was calculated. Subsequently, these values were normalized with the binned maximum background corrected intensity, which corresponds to the thickness where the reflected light constructively interferes. The average background corrected intensity of each bin is plotted as a function of ice thickness in blue, along with its standard deviation-based confidence interval (Fig. 5D).

As expected, for empty holes there is no intensity difference in the images before and after deposition, also corresponding to a thickness of zero. The maximum intensity at a thickness of around 90 nm corresponds to constructive interference of the light reflected at the air-water interfaces. At layer thicknesses larger than 90 nm, the intensities decrease down to negative values for destructive interference for thicknesses exceeding 180 nm. Negative values are the consequence of the fact that the carbon foil around these holes also holds a layer that destructively interferes. Therefore, there is no light diffracting into the holes making the intensity lower than that of the empty grid. Together, a continuous relationship reminiscent of one half of a sine wave dependency exists between image intensities and ice thicknesses from 0 – 180 nm.

#### 2.2.4. Optical model

In addition to the measured dependencies, the total amount of light reflected can also be calculated theoretically by thin film interferometry. The reflectance can be determined analytically using equation ^34^:

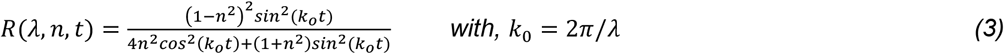

Where *R* is the reflectance, defined by the ratio between reflected power and incident power, *n* is the index of refraction, *k*_0_ is the wavenumber of light in free space, *λ* the wavelength of the light, and *t* is the layer thickness. Typical refractive index increments for protein solutions are around 0.2 ml/g ^35^, which would correspond to a change in refractive index of 0.004 for a protein concentration of 20 mg/mL. Therefore, the refractive index of pure water was used in this calculation.

The effective reflectance was determined by calculating the product of the components present in the light path of our setup. These are the incident light, applied filters, camera sensitivity and expected reflectance of the deposited sample. We calculated the effective reflectance as measured with the camera by integrating over the wavelength spectrum:

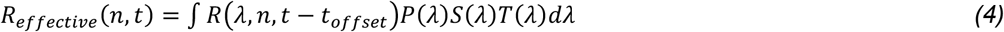

Here, *p*(*λ*) is the intensity of the incident light, *S*(*λ*) is the camera sensitivity and *T*(*λ*) the transmission of the applied long pass filter. The optical model was fitted with horizontal offset (*t*_*offset*_) of 17 nanometer matching the measured relationship in the plot. The theory and experimental data match up to an ice thickness of 180 nm. When inspecting the confidence interval of the scatterplot, derived from the standard deviation of the measurements, the accuracy can be put forward. In cases of thin ice (10 - 40 nm), measured intensities can predict the thickness with an error below ± 10 nm. At larger ice thicknesses up to 70 nm, the confidence interval is significantly broader reducing the accuracy to ± 20 nm. The optical model based on interferometry follows the relationship between optical intensity and ice thickness with good accuracy for relevant thicknesses used in single particle cryo-EM structure determination. This relationship can therefore be used as a calibration curve for reliable ice thickness estimation based on optical images.

### 2.3. Ice thickness control by pin printing

#### 2.3.1. Pin print – thickness correlation

To determine the reproducibility and the effect of the deposition parameters on ice thickness, we vitrified grid with different conditions in the VitroJet. In each session, 12 grids were prepared with 6 different conditions, such that each condition was duplicated. In the first three of these sessions, the velocity was decreased from 8 – 0.25 mm/s while halving the writing speed for each consecutive grid. In the second three sessions, the standoff was altered from 10 – 35 µm with fixed increments of 5 µm. Therefore, for each condition, 6 replicates were vitrified. These grids were loaded in the electron microscope, and the ice thickness for each individual hole was determined using the energy filter method. The final dataset contains data from 70 grids of 700,232 evaluated holes, where a histogram and violin plot for each condition was generated (Fig. 6).

**Figure 6:**
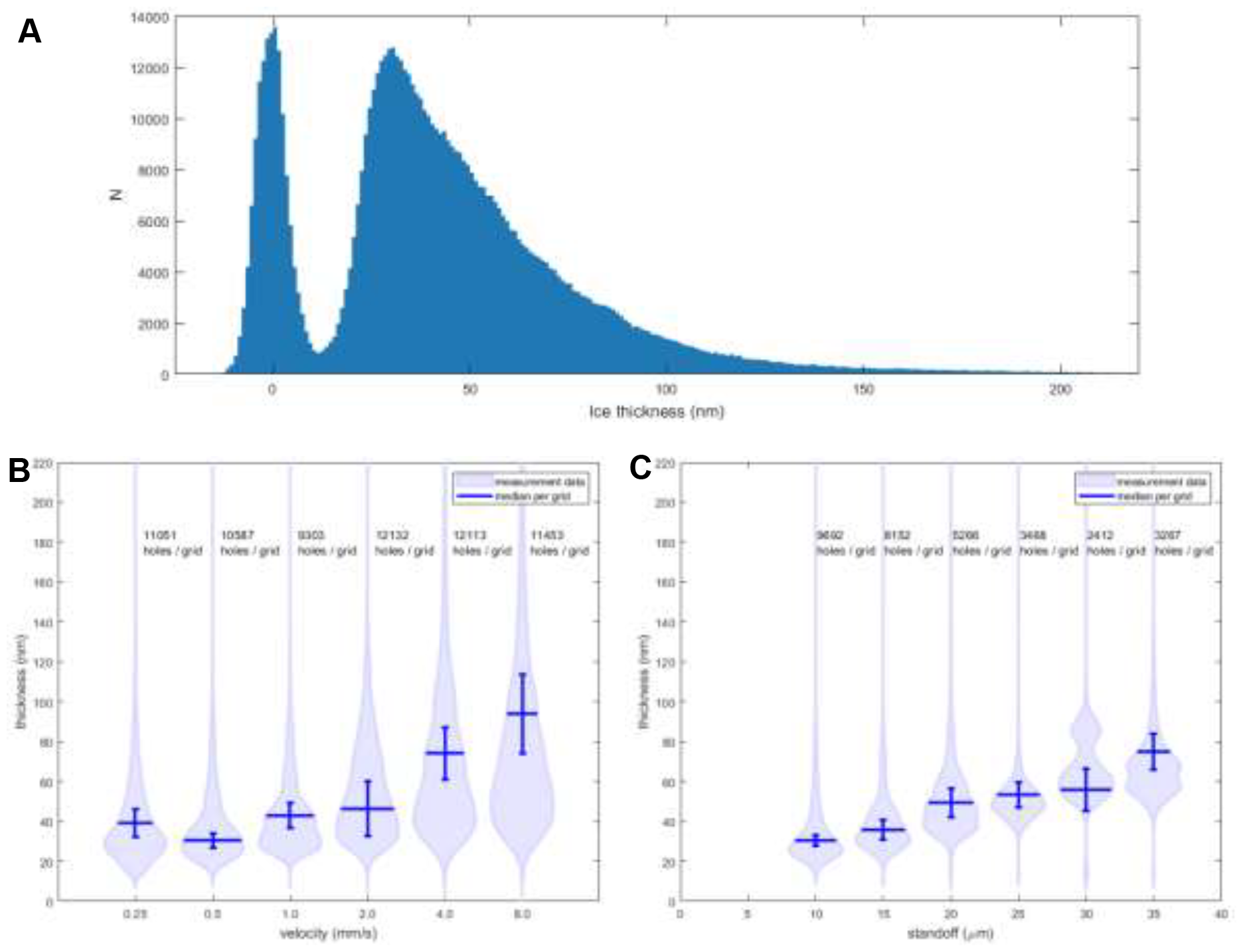
Relationship between pin print parameters in VitroJet and ice thickness measured in cryo-EM. (A) Histogram of the ice thicknesses in the dataset. (B and C) Violin plots of the ice thickness distribution when varying the writing velocity and standoff distance. The width of the violin corresponds with the relative occurrence of a certain thickness. The median ice thickness for each condition is shown in blue, and the standard deviation of the medians of the 6 replicates of this setting is shown with an error bar.

The histogram shows two peaks, where the first one corresponds to empty holes of which the maximum is shifted to a thickness of zero. The empty hole peak extends up to 12 nm, which is why only holes with a larger thickness are considered for further analysis. The second peak corresponds to the filled 559,749 holes, having a maximum at 30 nm and a tail extending up to 220 nm. The data per deposition setting is displayed in violin plots. For velocities up to 1 mm/s, the majority of holes are in the range of 20 – 50 nm. The median thickness for different grids can be reproduced with a standard deviation below ± 7 nm. When increasing the velocity from 0.5 to 8 mm/s, the median thickness and standard deviation increase steadily to 94 ± 20 nm, respectively. As a result, the range of available thicknesses for each condition increases. Thicker layers show a distribution within each square where the center is thicker and the holes around are thinner (Supplementary Materials A.2). The number of holes per grid that can be used for imaging in the microscope remains stable around 10,000 when altering the writing velocity.

When adjusting the standoff distance, the thickness changes steadily, and therefore the desired thickness can be tuned more easily. Increasing the standoff distance from 10 – 35 µm leads to a median thickness that varies from 30 – 75 nm with a standard deviation below ± 11 nm, whereas the number of filled holes reduces from approximately 10,000 to 3,000. Since the distance between pin and grid is larger, the liquid bridge of sample between them breaks up earlier and only part of the deposited area is covered. Additionally, the curvature of the liquid bridge is smaller, resulting in more wicking around the grid bars. This effect simultaneously raises the volume consumed per square and reduces the number of holes that are suitable for imaging in the electron microscope. The experiments demonstrate that tuning both writing velocity and the standoff distance results in effective coverage of the ice thickness ranges that are suitable for single particle electron microscopy.

#### 2.3.2. Theoretical model

A theoretical pin print model for has been developed that relates the deposition parameters to the expected ice layer thickness to predict the outcome from combinations of settings. A dimensional analysis is performed to analytically correlate the pin print parameters to the ice thickness (Supplementary Materials A.3). Dimensionless numbers are calculated to determine the relative importance of gravity, inertia, viscosity, and surface tension. This analysis shows that capillary forces are dominant under the pin. Additionally, the capillary forces are balanced by viscosity at the trailing edge of the pin where sample is deposited. Based on these relationships, the theoretical thickness can be determined:

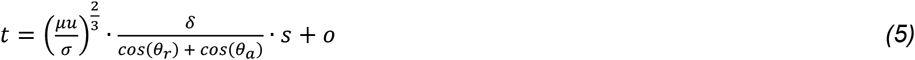

Where *t* is the thickness of the deposited layer, *µ* is the viscosity of the sample, *σ* is the surface tension of the sample, *u* is the writing velocity, *δ* is the standoff distance between pin and grid, *θ*_*r*_ is the receding contact angle at the pin, *θ*_*a*_ is the advancing contact angle on the grid.

The theoretical pin print model was fitted to the experimental data using the fluid properties of water and a respective receding and advancing contact angle of 0 ° and 30 °, resulting in a scaling factor *s* = 4.2 and offset *o* = 23 *nm* (Fig 7AB).

**Figure 7:**
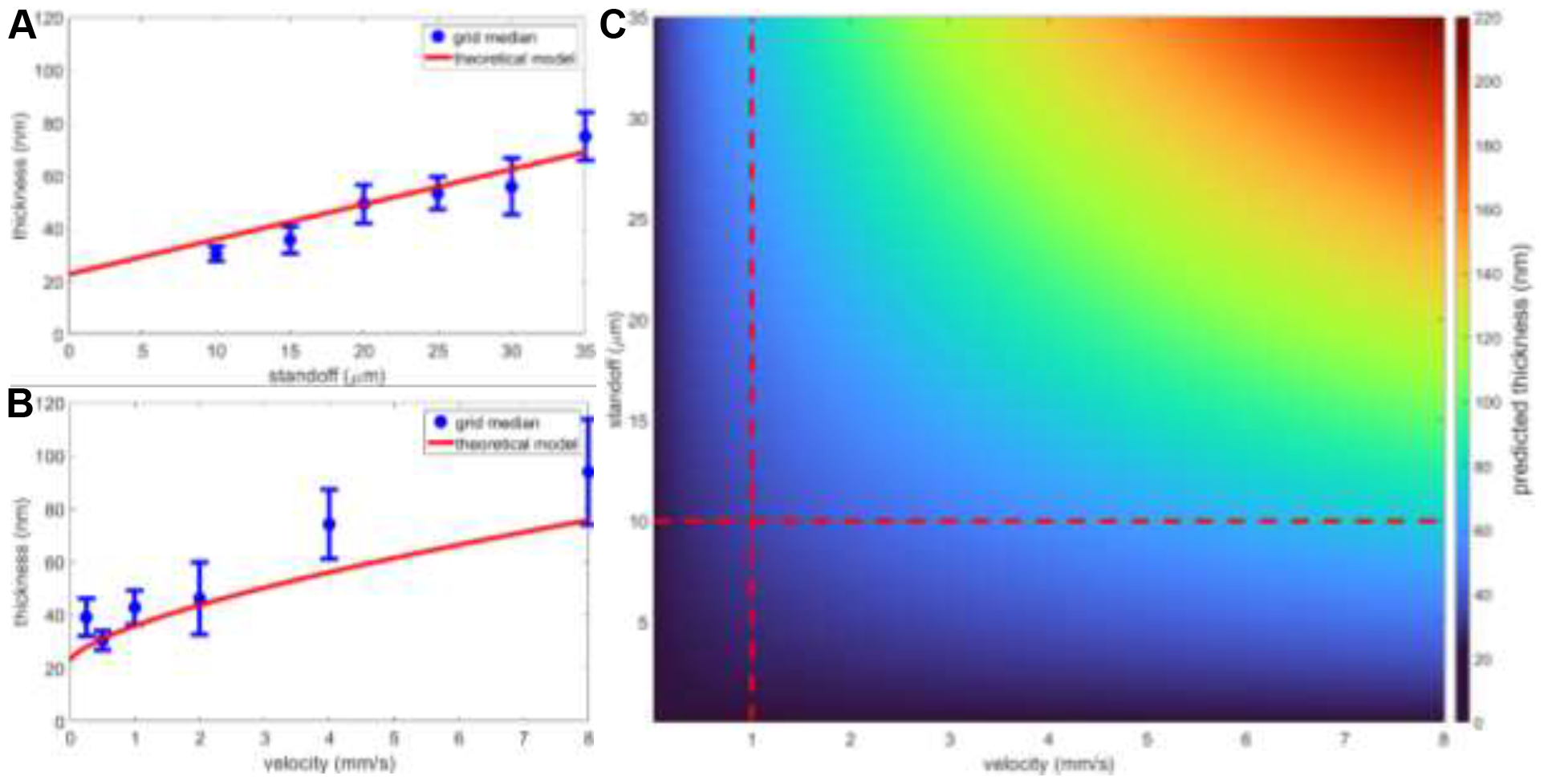
Correlation between the pin print data and theoretical model, and the predicted median ice thickness as a function of writing velocity and standoff distance. Relationship between ice thickness and pin printing parameters velocity (A) and standoff distance (B). The theoretical pin print model and measurement data confirm the correlation with a root mean squared error of 7 nm, where the ice is thinner at lower velocities and smaller standoff distances. (C) Estimated median ice thickness when using a combination of VitroJet protocol parameters, indicating the application space that can be utilized to tune desired ice thickness. The red dashed lines indicate where the model was validated with experiments.

The theoretical pin print model matches the experimental data and mostly remains within the measured standard deviation. The fit is a combination of the scaling using the theoretical pin print model and the influence of the grid itself. Therefore, we inspected grids from the same type and batch that were used in the experiments. These encompass an average grid bar width of 44 with a variation of ± 7 µm, which could influence hydrophilicity (n = 72, supplementary Materials A.4). Additionally, the carbon thickness was measured to be 59 ± 6 nm for this batch, which could directly affect ice thickness (n = 24 on 3 grids, supplementary Materials A.4).

Although there is variance in grids and sample preparation, the root mean squared error between measurement data and theoretical pin print model is 7 nm. The estimated median ice thickness as a function of writing velocity and standoff distance is displayed in a color plot (Fig. 7C). Experimental data was measured at the red dashed lines, yet a combination of settings can push the range of attainable ice thicknesses up towards 220 nm. When inserting sample properties in the theoretical relationship, parameters to obtain desirable ice thickness for specific projects can easily be determined.

### 2.4. Single particle structure determination

To demonstrate the relevance of tuning ice thickness for cryo-EM data collection, two datasets were acquired on different squares of the previously analyzed grid (See Fig. 2A, dashed blue squares labelled 1 and 2). The average ice thickness of square 1 was optically estimated according to the above established relationship at 30 nm, while square 2 had a thickness of 70 nm (26 nm and 67 nm from the energy filter method, respectively). As expected, the cryo-EM square overview and micrographs show higher contrast at the same defocus when the ice is thinner (Fig 8A-D). For both datasets, the micrographs were processed by motion correction and followed by contrast transfer function (CTF) estimation (Fig. 8E). The maximum resolutions obtained for fitting the CTF in the thin ice dataset is between 3 – 5 Å, whereas in the thick ice dataset fitting can only be successfully achieved between 5 – 7 Å resulting in poorer accuracy of CTF estimation. A total of 300,000 particles were selected for each dataset and included for a 3D reconstruction of apoferritin (Fig. 8FG). The resolution of the square 1 dataset with thin ice is better at 2.3 Å resolution over 2.5 Å resolution for square 2 with thick ice. When computing structures from subsets of the data and plotting the results in a Rosenthal-Henderson plot ^36^, the B-factor for both datasets were estimated at 168 and 240 Å^2^, respectively (Fig. 8H). Hence, when aiming for 3 Å resolution, only 3967 particles are needed when having thin ice, while using thick ice requires 14,502 particles to reach the same resolution. Effectively one needs 3.7 times less particles and thus microscope time to achieve the same result, in addition to reduced computational resources when analyzing smaller datasets.

**Figure 8:**
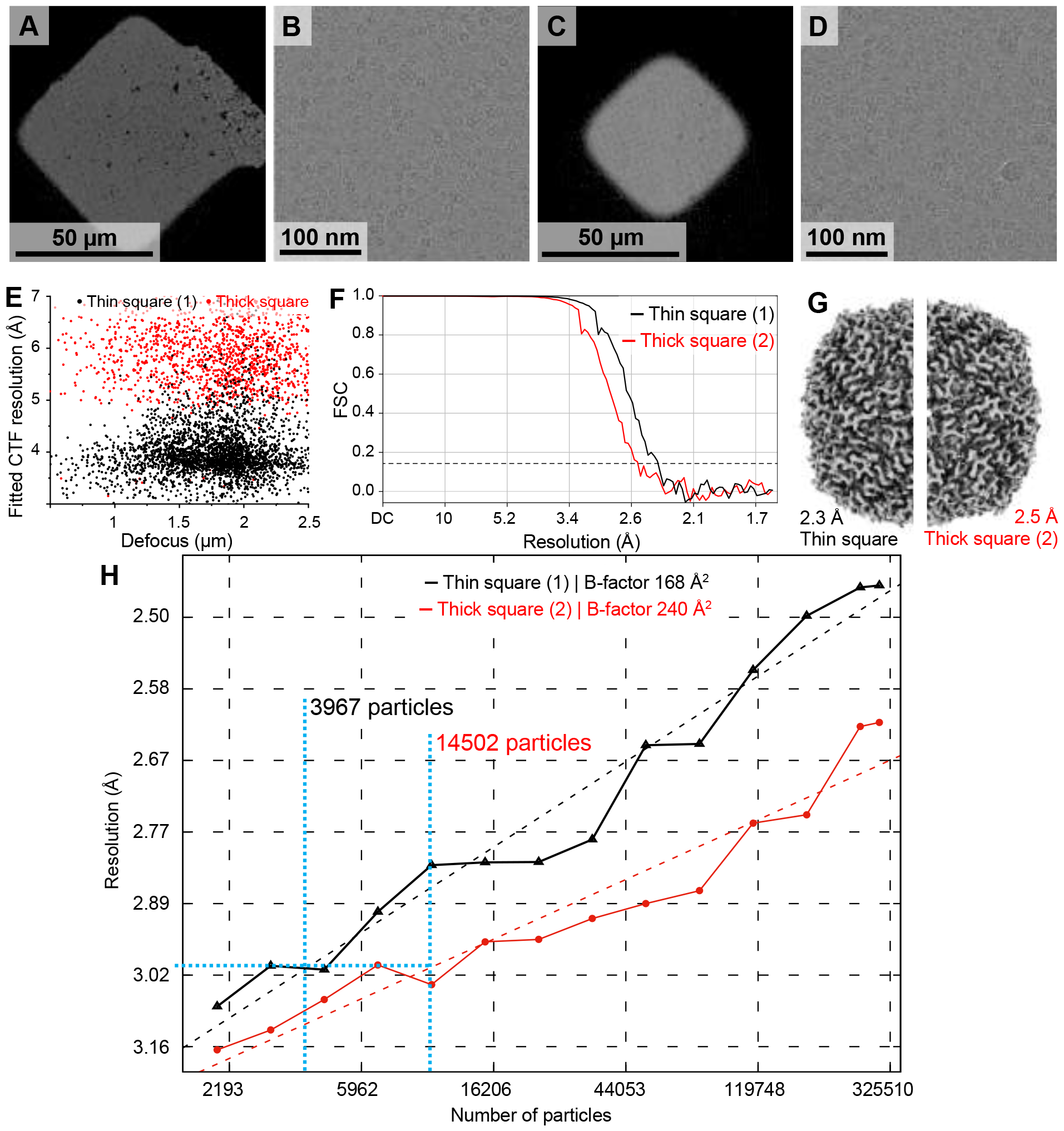
Comparison of two ice thicknesses and their influence on 3D structure determination of apoferritin. (A) Grid square 1 overview and (B) representative micrograph of apoferritin with thin ice of 30 nm. (C) Grid square 2 overview and (D) representative micrograph of apoferritin with thick ice of 70 nm. Representative micrographs were recorded at the same defocus. The locations of grid squares are marked on the atlas in Figure 3A in dashed blue boxes. (E) Scatterplot of fitted CTF resolution from square 1 (black) and square 2 (red) datasets. High ice thickness limits fitted CTF resolution for the dataset recorded on grid square 2. (F) Fourier shell correlation (FSC) of equally sized batches of 300,000 particles from grid square 1 (black) and 2 (red) selected for structure determination. (G) Reconstructed maps from square 1 (left) and 2 (right) show expected level of details, e.g., side chain densities, at 2.3 Å and 2.5, respectively. (H) Rosenthal-Henderson plot for square 1 (black) and square 2 (red) datasets. Determined B-factors were calculated at 168 Å^2^ and 240 Å^2^, respectively. To achieve 3 Å resolution, the thin ice sample requires 3.7 times fewer particles, resulting in remarkable reduction of data collection time.

## 3. Discussion

The results presented in this study demonstrate that the ice thickness per hole can be measured optically, and that the thickness distribution across the grid is controlled by the velocity and standoff distance. In single particle cryo-EM, the relevant ice thickness typically ranges between 10 – 100 nm. We show that images captured by the optical camera in the VitroJet can be analyzed based on hole intensities and correlated with ice thickness derived from cryo-EM images using the energy filter method (Figures 1 – 4). The optical layer thickness determination is theoretically based on thin film interferometry, and therefore suitable for predicting ice thicknesses up to the first constructive interference point. The ice thicknesses can be estimated in a range between 0 – 70 nm with an error below ± 20 nm, and with an error below ± 10 nm for the range between 10 - 40 nm (Figure 5). Outside this range, the ice thickness can still be estimated, but the accuracy decreases. Moreover, the ice thickness can be tuned resulting in a median thickness up to 75 nm using pin printing parameters and reproduced with an accuracy below ± 11 nm (Figure 6). Results match with the theoretical pin print model based on scaling of fluid dynamics forces, which can be used to optimize deposition parameters based on actual sample properties (Figure 7). Finally, on-the-fly tuning of ice thickness can be used to assess and select squares of desirable ice. Resulting resolution maps show a 3.7 x smaller number of particles required when comparing 30 and 70 nm ice thickness (Figure 8).

Thus far, the use of an electron microscope ^14–16^ or a dedicated optical light microscope or camera ^17–21^ was required to obtain feedback from sample preparation. In our study, the setup integrated in the VitroJet is used to measure ice thickness, eliminating the need to transfer grids and therefore avoiding potential grid damage and contamination. Additionally, previously presented optical methods provided a thickness prediction on square or atlas level ^17–19,21^, whereas our ice thickness measurement yields an estimation for each individual hole in the perforated foil of the grid. Adjusting blotting parameters was reported to not have a systematic effect on thickness ^23^, so that blotting does not allow for thickness tuning during grid preparation. For recently developed droplet-based or scribing methods, only qualitative relationships between deposition parameters and ice thickness were reported ^19,37^. Our results show that pin printing results in a reproducible distribution of ice thickness, enabling for selection of holes with optimal ice thickness in the electron microscope.

The accuracy of the optical measurement method is restricted by several factors. Due to the sinusoidal relationship between optical intensity and ice thickness, one optical intensity value can correspond to multiple ice thicknesses. Since ice thickness gradients are continuous, this ambiguity can be mitigated when consulting the intensities of surrounding holes to determine the actual thickness faithfully. Also, the optical model of layer thickness estimation by interferometry was given a negative horizontal offset to compensate for overestimation of ice thickness (Equation 4). This offset could either be due to the difference in liquid and solid phase during respective optical and electron microscopy, or since a negative correction factor was applied to the apparent mean free path obtained from reference values (Equation 2). Furthermore, the reflection of light depends on the index of refraction of the sample, which varies between different samples. However, most cryo-EM samples are in water-based buffers containing low concentrations of additives, in which the index of refraction will not significantly deviate and thus not influence the layer thickness estimation. Regarding sample deposition by pin printing, several aspects need to be considered as well. A startup transient at the beginning of the deposition pattern is observed, resulting in a thicker center and gradient towards the outside. Additionally, the reproducibility between grids decreases at higher velocities, potentially originating from a significant wicking from the thicker layer into the grid bars. Finally, both methods rely on grid geometry and materials. Batch-to-batch variability in grids could lead to deviations in hydrophilicity and foil thickness, affecting the final thickness. For reliable prediction of layer thickness from the optical images, it is important that the holes are sufficiently large for accurate intensity detection. It is expected that the background correction compensates at least partially for inaccuracies due to diffraction at smaller hole diameters. The flatness of the grid directly influences image sharpness and standoff distance, introducing a dependency on the operator skill when clipping grids. Pre-clipping at room temperature should help to simplify this procedure and decrease the number of bent grids.

Some of these concerns could be addressed by improving the setup. A monochromatic light source can be used in the optical setup such that only one wavelength contributes to the signal, simplifying the optical model and enabling selection of optimum accuracy at the thickness range of interest. Additionally, the algorithm to find holes and determine the intensity in holes could be further developed, incorporating details on hole pattern in the hole detection and averaging over multiple frames.

Since frames are recorded from start to end of the deposition, it is possible to monitor the evolution of the layer over time. Using this information, dewpoint settings can be tuned to minimize evaporation or condensation. Furthermore, the fluid flow and stability of the sample on the grid during and after deposition can be studied. Controlling ice thickness with pin-printing can still be enhanced in the future. Developments in new grid types and supports based on MEMS technology may improve accuracy and reproducibility ^38^. In case these grids are still curved or tilted, a height map could be recorded with the integrated camera. The path of the pin can be adjusted to compensate for this curvature, maintaining a constant standoff distance. The presented experimental data demonstrates how tuning of a single parameter can be used to tune ice thickness; however, a combination of multiple parameters can be used to enhance effectivity and provide flexibility for compensation for sample fluid properties.

The results show that the ice thickness can be tuned when using the pin-print deposition technique and optical measurement. Using the camera, the layer thickness in individual holes of EM-grids can be determined optically. Pin-printing provides reproducible sample deposition and enables tuning of the ice thickness distribution. The accuracy of the proposed methods is sufficient to allow for on-the-fly measurement and control of deposition to achieve optimal layer thicknesses without the use of an electron microscope. As shown for the test sample apoferritin, this can decrease time to target holes with suitable ice thicknesses during screening. Recording an atlas on a grid takes approximately 10 min, for an autoloader system with 12 grids this results in 2 full hours. If the desired thickness is known and the optical thickness maps are available, this could substantially decrease the number and size of the recorded atlases, and allow targeting the most promising grids and holes directly.

## 4. Materials and methods

### 4.1. Grids and sample

Copper 200 mesh, carbon R2/1 foil grids (Quantifoil Micro Tools GmbH, Germany) were pre-clipped into an autogrid ring (Thermo Fisher Scientific Inc, MA, USA) at room temperature. Before use, the autogrids were glow discharged in a PELCO easiGlow (Ted Pella Inc., CA, USA) for 60 s at 15 mA, 0.4 mbar including a hold time of 10 seconds. The sample apoferritin from horse spleen was purchased (Sigma-Aldrich / Merck KGaA, Germany), buffer exchanged and concentrated using Amicon Ultra filtration units (Merck KGaA, Germany) to 20 mg/mL in 5 mM Hepes, pH 7.5 and 25 mM NaCl. Aliquots were flash frozen and stored at a temperature of −80 °C before use.

### 4.2. Grid preparation

To determine the correlation between the optical camera images and ice thickness, the VitroJet (CryoSol-World B.V., The Netherlands) was used to prepare and vitrify grids. Inside the VitroJet, the autogrids were treated using the integrated plasma cleaner to ensure reproducible hydrophilicity between grids. Afterwards, the sample was deposited onto the autogrids using pin printing. In this process, a solid tungsten pin dipped in the pipette to pick up approximately 0.5 nL. Afterwards, the pin approached the foil of the grid to a set standoff distance, such that the sample forms a bridge between pin and grid. A thin layer was deposited by moving the pin laterally to the grid with a predefined velocity. Deposition started in the center and spiraled out to cover a circle with a diameter of 800 µm. Sample deposition occurred in a climate chamber. During deposition, both pin and grid were controlled to dewpoint temperature to minimize evaporation. After deposition, the autogrid was vitrified by jet-vitrification.

Sample deposition was monitored by a camera that is integrated in the VitroJet (Fig. 1). The setup had on axis illumination (white LED, SL112, Advanced Illuminations, Vermont, USA) with an absorptive filter and a 495 nm long pass filter (NE20B and FGL495, respectively, Thorlabs GmbH, New Jersey, USA). Light reflected from the grid was captured using a microscope objective (M Plan Apo NIR, 20X, Mitutoyo, USA) and recorded by a camera (acA2440-20gm, 5MPix, Basler AG, Germany). The images were recorded in real time at a frame rate of 20 fps and stored on a personal computer (HP, USA).

#### 4.2.1. Protocols used for optical correlation

A total of 18 grids were prepared using the VitroJet and subsequently imaged both by the optical camera and in cryo-EM. All grids were plasma cleaned for 60 s, had a deposition temperature of 4 °C and a pin and grid dewpoint offset calibrated to 0.1 °C and −0.3 °C, respectively. A single application with a circular spiral pattern was used. To create a wide range of layer thicknesses, several protocols with distinct deposition parameters were used. With a standoff of 10 µm and a spacing of 150 µm, 6 grids were made with a velocity of 0.5 mm/s and 3 grids for velocities 1 and 2 mm/s. With a standoff of 15 µm, 3 grids were made with a spacing of 150 µm and velocity 1 mm/s, and another 3 grids with spacing 70 µm and velocity 5 mm/s.

#### 4.2.2. Protocols used for pin-print correlation

To determine reproducibility and controllability with pin-printing, a total of 72 grids were prepared in 6 experiments with the VitroJet. For all grids, a plasma cleaning duration of 60 seconds was selected. Grids were vitrified with a deposition temperature of 4 °C and a pin and grid dewpoint offset calibrated to −0.2 °C and −0.6 °C, respectively. A single application with a circular spiral pattern having spacing between subsequent spirals of 70 µm was used.

In the first half of these sessions, the standoff was fixed at 10 µm and the velocity was varied between 0.25 – 8 mm/s, doubling the writing speed between different grids. In the second half of the experiments, the velocity was set at 1 mm/s and the standoff was altered from 10 – 35 µm with fixed steps of 5 µm. Two grids were lost during processing, resulting in a final dataset of 70 grids.

### 4.3. Cryo-EM analysis

#### 4.3.1. Ice thickness measurement for optical correlation

The ice thickness was measured using a Talos Arctica (Thermo Fisher Scientific Inc, MA, USA) operated at 200 kV and equipped with a Gatan K3 camera behind a post-column Bioquantum energy filter (Gatan Inc, CA, USA). In this case, an apparent mean free path of 485 nm is used according to the acceleration voltage of 200 kV and medium magnification during imaging ^15^. The correction factor is determined by shifting the peak of the empty holes in the histogram to zero by 9 nanometer.

Atlas images were acquired at 110 x in LM mode, whereas for recording grid square images, the magnification LM 320 x with a pixel size of 26 nm was chosen. These settings ensured that each image showed exactly one grid square in the field of view. In addition, the intensity was set such that the grid square will receive the final dose of 105 e/pix/s. For those the grid squares that were analyzed with the optical analysis, the eucenteric height was set and an unfiltered image of the grid square was recorded. Afterwards, the energy filter with a slit width of 20 eV was inserted and a filtered image was recorded, where the zero-loss peak manually aligned at the LM magnification.

For each square image in absence and presence of the energy filter, the region close to the grid bars of the square containing thick ice was excluded from further analysis using intensity thresholding. From the remaining area of the square, the expected number of holes was calculated based on the known grid properties of hole diameter and spacing. Circles with the expected diameter were detected using the Hough transform ^33^. Subsequently, false positives were discarded based on implausible distances to nearest neighbors and apparent ice contamination. The average intensity for each hole was determined from a circular area with 85% of the hole diameter to exclude any carbon edges.

#### 4.3.2. Ice thickness measurement for pin-print correlation

For this analysis, a Titan Krios G4 was used equipped with a C-FEG, SelectrisX energy filter and Falcon 4i detector, operated at 300 kV. To calibrate the mean free path for this microscope, a total of 43 tilt series were acquired from one grid. Data was recorded using a dose symmetric tilt scheme with a tilt range of +/- 60°, tilt increment of 20° (61 total tilts) and a group size of 4 at a nominal magnification of 42,000x (3.033 Å calibrated pixel size), a total dose of ~120 e-/Å2 and a 10 eV energy filter slit width ^39^. The tilt series were constructed into tomograms using AreTomo with a binning of 4 and a z-height of 256 pixels in the final volume ^40^. In the tomograms, the ice thickness was determined manually by locating the ice sheet in the cross-section of each tomogram. A correction factor was used to shift the empty hole peak to zero ^15^. Afterwards, the apparent mean free path was determined by a linear fit of ln(I_tot_ / I_ZLP_) against the measured thicknesses from the tomograms ^14^. This resulted in a mean free path of 352 nm and a correction factor of 10 nm.

To determine the ice thickness on a large number of grids, square maps without and with the energy filter slit inserted were acquired in a semi-automated manner using SerialEM on the same Titan Krios microscope ^41^. First, an atlas overview of up to 12 grids loaded in the autoloader was acquired. Then, the centers of all grid squares from all grids selected for ice thickness measurements were manually selected and marked as acquisition points. A virtual map for each selected square was generated using the pyem SerialEM plugin with a target magnification of 470x (nominal, LM mode) ^42^. These maps were used for recentering the respective square upon reloading of the respective grid from the autoloader to the stage. After recentering, a 3 x 3 montage was acquired at 1,950x magnification (nominal, LM mode) using image shift, each without and with the slit of the energy filter inserted (ZLP, 30 eV slit width). Once all selected squares from a grid were acquired, the next grid was loaded onto the stage automatically and square map acquisition continued accordingly.

The grid overviews were processed by first combining the tiled raw data using IMOD’s “blendmont” function ^43^. The “imodholefinder” function in IMOD was utilized to detect hole positions in each of the resulting square maps. The mean value for each hole was determined from all pixels that were located within a circle (128 pixels in diameter) around the hole center (the entire diameter of a hole was ~330 pixels). Thus, ~15 % of all pixels from the entire hole contributed to the measurement, while the influence of a non-homogeneous ice thickness throughout the holes was mitigated. The measure consequently describes the intensity at the center of each hole.

#### 4.3.3. Single particle structure determination

Data was acquired on a grid prepared for the optical correlation with a writing velocity of 1 mm/s, standoff of 10 µm, and spacing of 70 µm. Single particle data was recorded on a Titan Krios G4 operated at 300 kV with a non-energy filtered Falcon-IV direct electron detector using EPU software v3.40 (Thermo Fisher Scientific Inc, MA, USA). In the two grid squares with thin and thick ice, 3780 micrographs and 1469 micrographs were recorded, respectively, at a nominal magnification of 96,000x with a calibrated pixel size of 0.808 Å and a total fluence of 30 e−/ Å^2^ in EER mode. A total of 7 exposures per 2 µm hole were acquired with faster beam-tilt acquisition in aberration-free image shift mode with a set defocus between −1.0 and −2.2 µm resulting in approximately 500 images/h. Data processing was performed in CryoSPARC v4.2.1. After patch motion correction with EER upsampling on a 30-frame grid and patch CTF estimation, exposures with a CTF fit worse than 5 Å were excluded for the thin ice square. Particles were picked with ring-shaped blob templates with a 60 Å inner and 110 Å outer diameter and extracted in 256 x 256 pixel squares. These particles were subjected to reference-free 2D classification and classes showing clear secondary structure were selected for further analysis. This subset was used for ab initio 3D map generation with a resolution range of 6 - 9 Å with elevated initial and final batch sizes of 300 and 1000, respectively. Ab initio models were low pass filtered to 12 Å prior to homogeneous 3D refinement with each 300,000 particles with O-symmetry imposed, including local and global CTF refinement (tilt and trefoil), followed by a local non-uniform refinement. Nominal resolutions at FSC = 0.143 for random particle subsets were computed using ResLog while the final B-factor calculation was performed by linear curve fitting in Gnuplot v6.0. Figures were generated using ChimeraX v1.4 and Adobe Illustrator 2023.

## 5. Acknowledgements

We are grateful to Giulia Weissenberger her contribution in developing and testing of these ideas, and Robin Loose for the image analysis algorithm for the electron microscopy images. We would like to thank Michiel Peters, Ben Bormans and Bob Ashley for critically reading the manuscript and their helpful feedback.

This research received funding from the Netherlands Organization for Scientific Research (NWO) in the framework of the take-off number [17007]. The authors gratefully acknowledge computing time on the supercomputer JURECA [1] at Forschungszentrum Jülich. The authors acknowledge the access and services provided by the Imaging Centre at the European Molecular Biology Laboratory (EMBL IC), generously supported by the Boehringer Ingelheim Foundation.

[1] Jülich Supercomputing Centre. (2021). JURECA: Data Centric and Booster Modules implementing the Modular Supercomputing Architecture at Jülich Supercomputing Centre Journal of large-scale research facilities, 7, A182. http://dx.doi.org/10.17815/jlsrf-7-182

