## Supplementary materials for "Ice thickness control and measurement in the VitroJet for time-efficient single particle structure determination"

### Appendix A

**Supplementary Materials**

#### A.1 Mean free path calibration

For the pin printing experiments, the apparent mean free path of the electron microscope is calibrated. The correction factor is determined by shifting the peak of the empty holes in the histogram to zero resulting in a correction factor 10 nm. After determining the correction factor for the pin print correlation, the mean free path for the 300 kV microscope was calibrated to 352 nm using tomography (Fig. A.1). This resulted in a root mean square deviation of 6 nm for the energy filter method to determine the ice thickness.

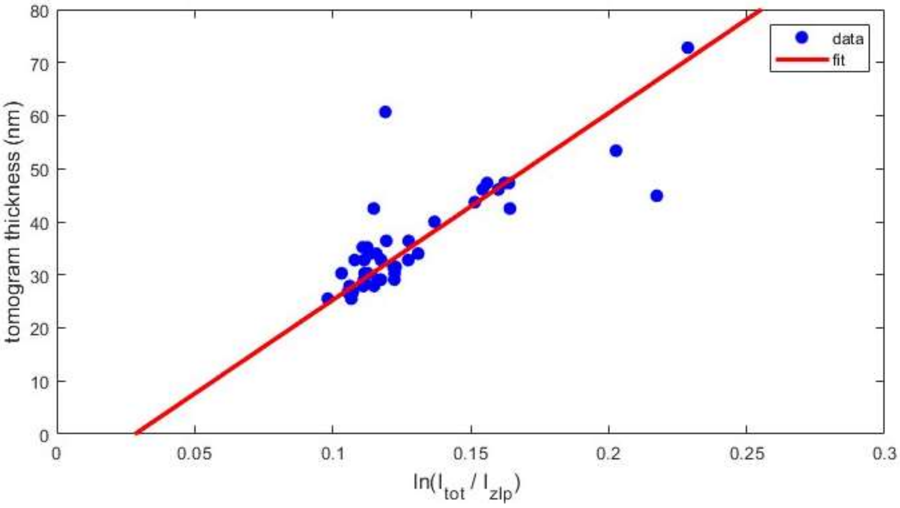


*Supplementary figure 1: Apparent mean free path calibration for the pin printing experiments. The natural logarithm of the intensity ratio in presence and absence of the energy filter was plotted with respect to the ice thickness measured by tomography. Using the correction factor and a linear fit, the apparent mean free path was determined to be 352 nm.*

#### A.2 Thickness distribution within squares

When depositing a thin layer, the intensity within the squares is uniform (Fig. A.2A). If a layer is deposited with a thickness that surpasses the maximum reflection at 90 nm, Newton interference rings start to form. These layers show a thickness distribution within each square, where both the center and the area close to the grid bars are thicker compared to the region in between (Fig. A.2B).

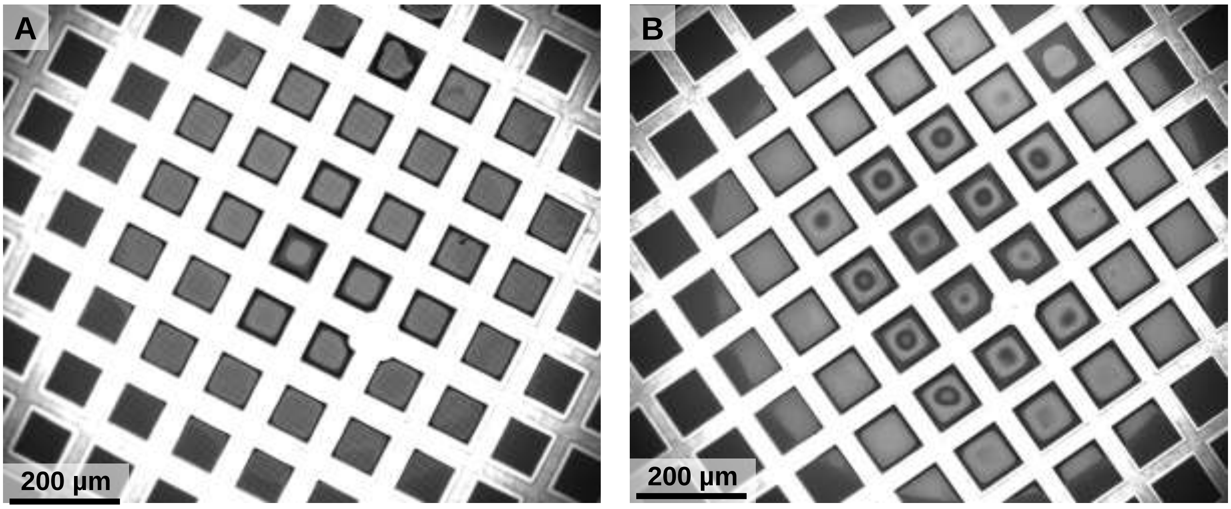


*Supplementary figure 2: Thickness distribution on a square with a thin (A) and thick (B) layer. The thin layer shows a uniform thickness in the squares, whereas the thicker layer contains a thickness distribution within each square. The center and area close to the grid bars are thicker with respect to the region in between.*

#### A.3 Theoretical model

A theoretical model has been developed that relates the deposition parameters to the expected ice layer thickness to predict the outcome from combinations of settings. A dimensional analysis is performed to analytically correlate the pin print parameters to the ice thickness. Since the aspect ratio between pin diameter and standoff distance is large, the situation is assumed to be 2 dimensional (Fig. A.3).

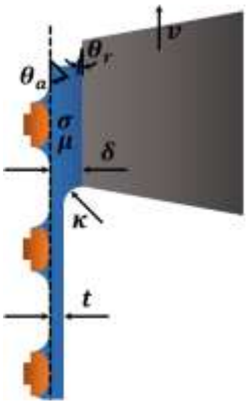


*Supplementary figure 3: Schematic overview of the parameters involved in pin printing, where a solid pin moves with respect to the grid to deposit sample. The thickness of the layer is dependent on writing velocity, standoff distance between pin and grid, hydrophilicity of pin and grid, and sample properties.*

First, the significance of gravitational forces over capillary forces is calculated using the dimensionless Bond number $Bo = \Delta\rho g L^2/\sigma = 7.8 \cdot 10^{-4}$, indicating that gravity is negligible. The importance of inertia compared to surface tension can be determined using the dimensionless Weber number $We = \rho u^2 \delta/\sigma = 1.4 \cdot 10^{-7}$, demonstrating that this situation is dominated by surface tension. Viscous forces can be scaled with the surface tension by the modified capillary number $Ca_D = \mu u D/\sigma\delta = 7 \cdot 10^{-5}$, showing that capillary forces are much larger than viscous friction under the pin. Therefore, the curvature of the meniscus at the front and back of the pin is equal. The curvature at the front of the pin is determined by the contact angles and standoff distance.

$$\kappa = \frac{cos(\theta_r)+cos\ (\theta_a)}{\delta} \qquad (A.1)$$

Where $\kappa$ is the meniscus curvature, $\theta_r$ is the receding contact angle at the pin, $\theta_a$ is the advancing contact angle on the grid, and $\delta$ is the standoff distance between pin and grid.

The importance of viscosity compared to inertia can be estimated with the Reynolds number $Re = \rho u\delta/\mu = 0.01$, showing that inertia is negligible compared to viscous forces at the pin printing velocities that were used. At the trailing edge at the back of the pin the layer thickness is much smaller, making the relative importance of inertia even smaller and that of viscous friction even larger. In the layer thickness that is deposited, capillary forces are balanced by viscosity.

$$\frac{\kappa\sigma}{L} \propto \frac{\mu u}{t^2} \qquad (A.2)$$

$$\frac{t}{L^2} \propto \kappa \qquad (A.3)$$

Where $\sigma$ is the surface tension of the sample, $\mu$ is the viscosity of the sample, $u$ is the writing velocity, $L$ the length of the dynamical meniscus, and $t$ is the thickness of the deposited layer. We solve these equations to determine the deposited sample thickness on the carbon foil.

$$t \propto \left(\frac{\mu u}{\sigma}\right)^{\frac{2}{3}} \cdot \frac{\delta}{cos(\theta_r) + cos(\theta_a)} \propto Ca^{\frac{2}{3}}\kappa^{-1} \qquad (A.4)$$

Afterwards, the theoretical model is fitted with the experimental data using a dimensionless scaling factor and add an offset to account for grid hydrophilicity and the thickness of the carbon foil itself.

$$t = Ca^{\frac{2}{3}}\kappa^{-1} \cdot s + o \qquad (A.5)$$

The scaling factor and the offset are determined with least squares fitting.

##### A.4 Grid inspection

To determine the origin of the variation in layer thickness and relative importance of various factors, the grids that were used in the pin-printing experiments were inspected in more detail. The optical camera of the VitroJet provides an overview of the grids before deposition, revealing variations on grid bar width of 44 ± 7 µm over 72 grids from the same batch (Fig. A.4AB). The thickness of the carbon foil was measured in electron microscope images of the edge of empty holes outside the deposition region with a stage tilt of 45 °, leading to 59 ± 6 nm on 24 holes of 3 grids (Fig. A.4CD).

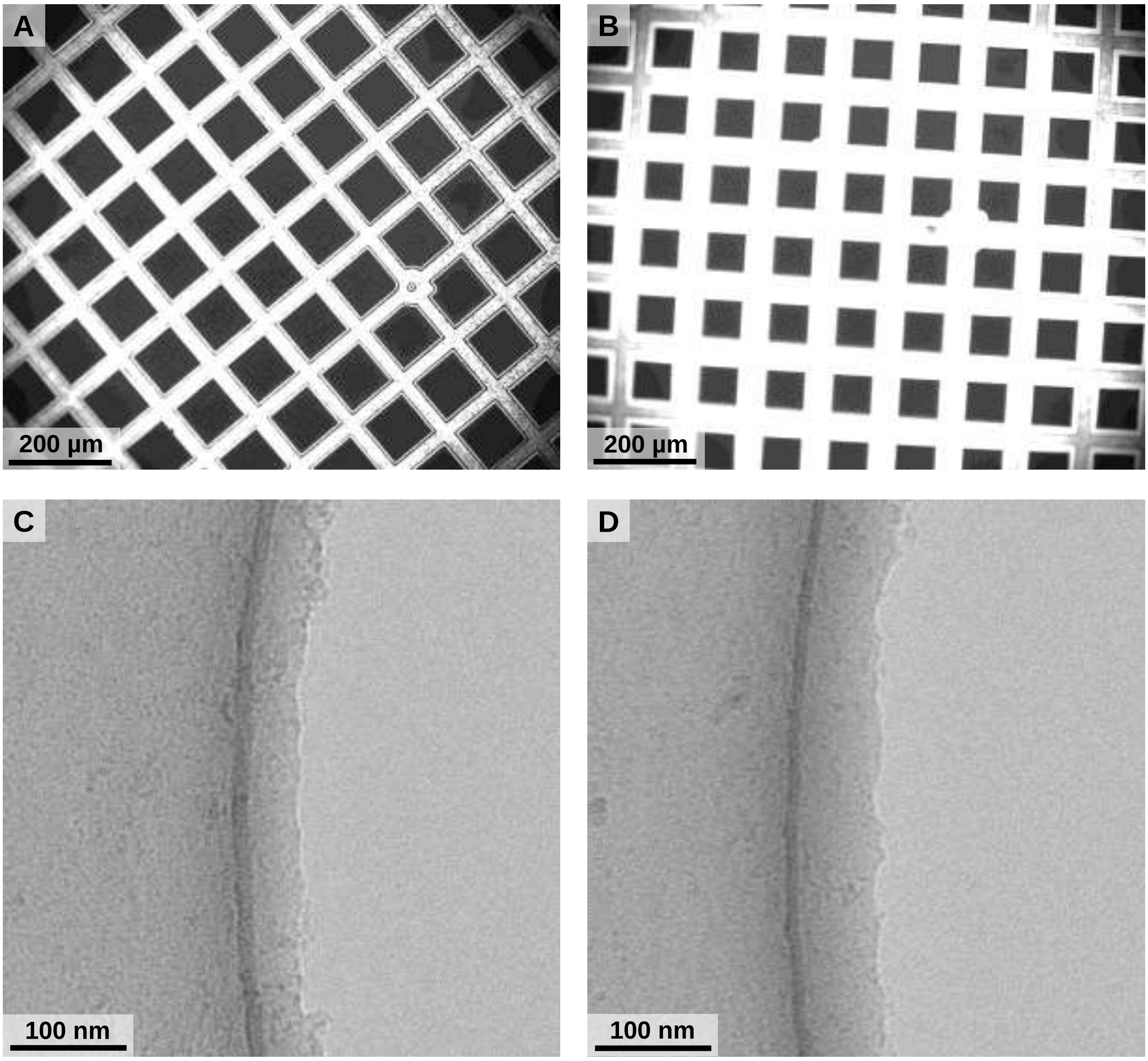


*Supplementary figure 4: Images of grids, (AB) optical images from the VitroJet camera showing variation in grid bar width. Electron microscopy images showing the edge of an empty hole with a stage tilt of 45°, indicating the thickness of the perforated carbon foil.*
